## Supplement for "Ciliary neuropeptidergic signaling dynamically regulates excitatory synapses in postnatal neocortical pyramidal neurons"

**SUPPLEMENTAL FIGURE LEGENDS**

**Figure 1 – figure supplement 1.** Gross neuronal morphology is unaltered upon acute knockdown of ciliary proteins.

**A,B)** Relative fluorescence intensities of immunolabeled ciliary ARL13b **(A)**, and cilia lengths **(B),** in neurons transfected with the indicated plasmids 48 hrs post-transfection. Each dot is a measurement from a single neuron. Values in **I** are normalized to values in controls. Bars are average ± SEM. ** and *** indicate P<0.01 and 0.001 for the indicated conditions (Wilcoxon rank-sum). n: Control = 19, shARL13b_2 = 26; 3 dissociations.

**C)** Representative images of DIV11 pyramidal neurons expressing the indicated plasmids 48 hrs post-transfection. Scale bar: 50 µm.

**D, E)** Total lengths **(D)** and branch points **(E)** of apical-like dendritic arbors of cultured pyramidal neurons expressing the indicated constructs at 48 hrs post transfection. Each dot is a measurement from a single neuron. Bars are average ± SEM. n: Control = 21, shArl13b_2 = 14, shIft88/shCep164 =17; 3 dissociations.

**F, G)** Relative fluorescence intensities of immunolabeled ciliary IFT88 **(F)**, and cilia lengths **(G),** in neurons transfected with the indicated plasmids 24 or 48 hrs post-transfection. Each dot is a measurement from a single neuron. Values in **F** are normalized to values in controls. Bars are average ± SEM. *, ** and *** indicates P<0.05, P<0.01,  P<0.001, respectively for the indicated conditions. n: (24 h) n: Control = 23, shIft88/shCep164/GFP = 25; 3 dissociations; (48 h) Control = 43, shCep164/GFP = 20; shIft88/GFP = 17, shIft88/shCep164/GFP = 23; 3 dissociations.

**H)** Representative images of neurons expressing GFP alone (top) or shIft88/shCep164 and GFP (bottom). Cilia were immunolabeled with antibodies against AC3 and IFT88. Images at right show enlarged (2.5X) views of cilia (yellow boxes). Scale bars: 5 µm.

**Figure 2 – figure supplement 1.** Presynaptic VGlut1 staining is increased upon acute knockdown of ciliary proteins.

**A, B)** Relative fluorescence intensity of immunolabeled VGlut1 **(A)** or GAD65 **(B)** at GluA2/VGlut1 or Gephyrin/GAD65 colocalized puncta, respectively, in neurons transfected with the indicated plasmids at 24 hrs or 48 hrs post transfection. Intensity values are normalized to values in controls. Each dot is the average summed pixel value of examined synapses per neuron. Bars are average ± SEM. * indicates $P<0.05$ for the indicated conditions; additional P-values are also indicated. **(A)** n: (24 hrs) Control = 25, shArl13b_1 = 24; (48 hrs) Control = 32, shArl13b_2 = 19, shIft88/shCep164/GFP = 39; 4 dissociations. **(B)** n: (24 hrs) Control = 17, shArl13b_1 = 22 neurons; 4 dissociations; (48 hrs) Control = 25, shArl13b_2 = 19, shIft88/shCep164/GFP = 23; 3 dissociations.

**Figure 5 – figure supplement 1.** SSTR3 is present in the majority of the cilia of NeuN-positive neurons and absent from multiple inhibitory neuron subtypes in the cortex.

**A)** Representative images of NeuN, GAD67 and SSTR3 co-labeling in fixed cortical tissue stained. Center and right panels are enlarged representative images from each indicated layer of cortex. GAD67+ neuronal soma are outlined in dashes, arrows point to cilia. Scale bars: 50 μm (left panel) and 5 μm (center and right panels).

**B)** Quantification of the percentage of neurons positive for NeuN and SSTR3 (top), and GAD67 and SSTR3 (bottom) by layer of cortex. n: NeuN+ = 1,520, GAD67+ = 253; 6 animals.

**C)** Representative images of inhibitory subtypes of neurons in fixed cortical tissue stained with DAPI. Neurons were immunolabeled with antibodies against indicated subtype specific markers and either PCTN and SSTR3, or with AC3, PCTN and SSTR3. PCTN was used to identify centrioles and basal bodies (arrows) Scale bar: 5 µm.

**Figure 6 – figure supplement 1.** Agonism and antagonism of SSTR3 alters excitatory synaptic strengths.

**A)** Relative fluorescence intensity of VGlut1 at colocalized Shk3/VGlut1 puncta on neurons treated with 2 µM SSTR3 agonist L-796,798 or 1 µM MK-4256 SSTR3 antagonist. Cultures were immunostained at the shown times following treatment. Intensity values are normalized to values in control neurons. Each dot is the average summed pixel value of all examined synapses per neuron. Bars are average ± SEM. * and ** indicate $P<0.05$ and 0.01, respectively for the indicated conditions; (LMM with Dunnett-type correction for multiple comparisons); additional P-values are also shown. n: (6 hrs) Control =23, agonist = 10, antagonist = 17; (18 hrs) Control = 35, agonist = 33, antagonist = 17; (24 hrs) Control = 87, agonist = 17, antagonist =40; ≥3 dissociations.

**B)** Relative fluorescence intensities of immunolabeled Shk3 and VGlut1 at colocalized Shk3/VGlut1 puncta. Neurons were treated with the indicated concentrations of the SSTR3 agonist L-796,778 or antagonist MK-4256 for 24 hrs. Each dot is the average summed pixel value of examined synapses per neuron. Bars are average ± SEM. *, ** and *** indicate $P<0.05$, $P<0.01$ and 0.001, respectively for the indicated conditions; (LMM with Dunnett-type correction for multiple comparisons). n: Control = 87, 0.5 µM agonist= 26, 1 µM agonist = 25, 2 µM agonist= 17, 0.125 µM antagonist = 19, 0.5 µM antagonist = 15, 1 µM antagonist = 40; ≥3 dissociations.

**C)** Number of colocalized Shk3/VGlut1 puncta per μm of dendrite analyzed (density) on neurons treated with the indicated concentrations of the SSTR3 agonist L-796,778 or antagonist MK-4256 for 24 hrs . Each dot is the density of synapses examined per neuron. Bars are average ± SEM. * indicates $P<0.05$ for the indicated conditions; (LMM with Dunnett-type correction for multiple comparisons). n: As in **B**.

**D)** Lengths of cilia in neurons treated with SSTR3 agonist and antagonist 24 hrs following treatment. Each dot represents a measurement from a single neuron. Bars are average ± SEM. n: Control = 63, 2 μM agonist= 7, 1 μM antagonist = 33, ≥3 dissociations.

**E)** Fraction of alive and dead neurons at 24 hrs following treatment with SSTR3 agonist or antagonist. Neurons were co-stained with propidium iodide and Zombie Green; cells positive for either or both stain were scored as dead. n: Control = 97, 2 μM agonist = 92, 2 μM antagonist = 33; 3 dissociations.

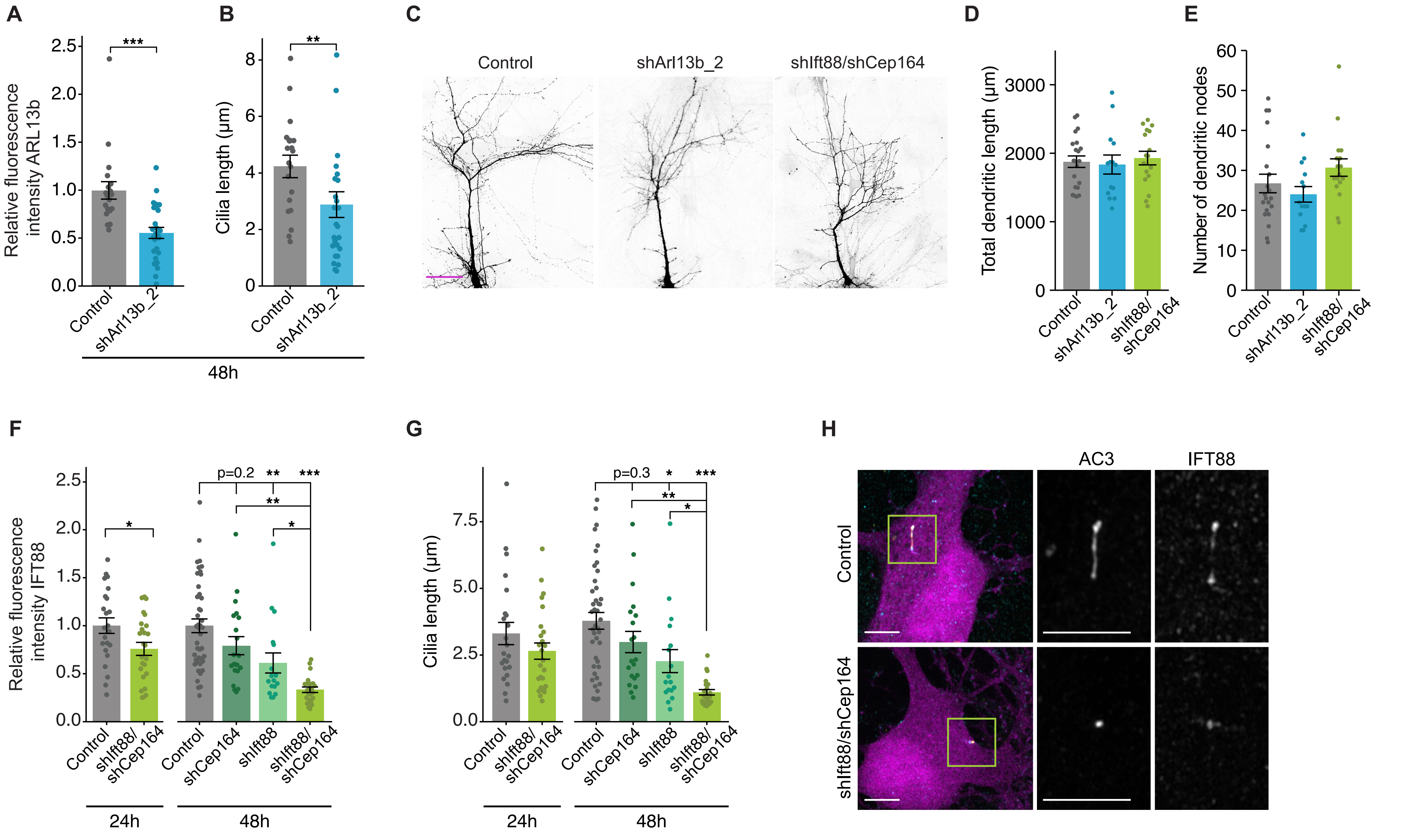


Tereshko et al.,
Figure 1 - figure supplement 1

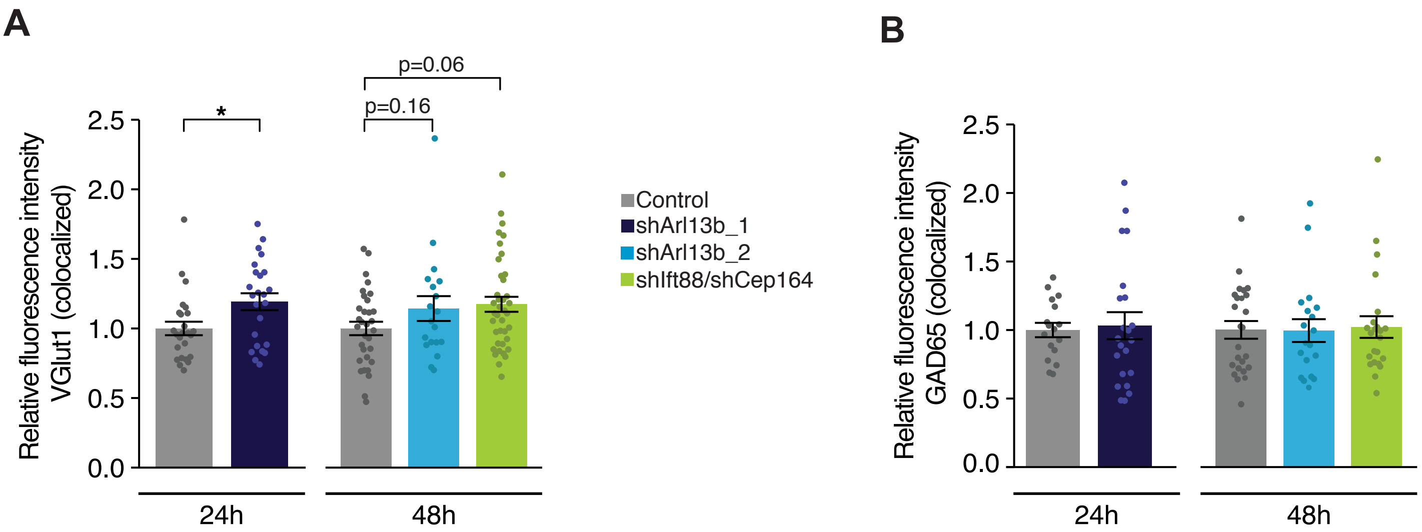


Tereshko et al.,
Figure 2 - figure supplement 1

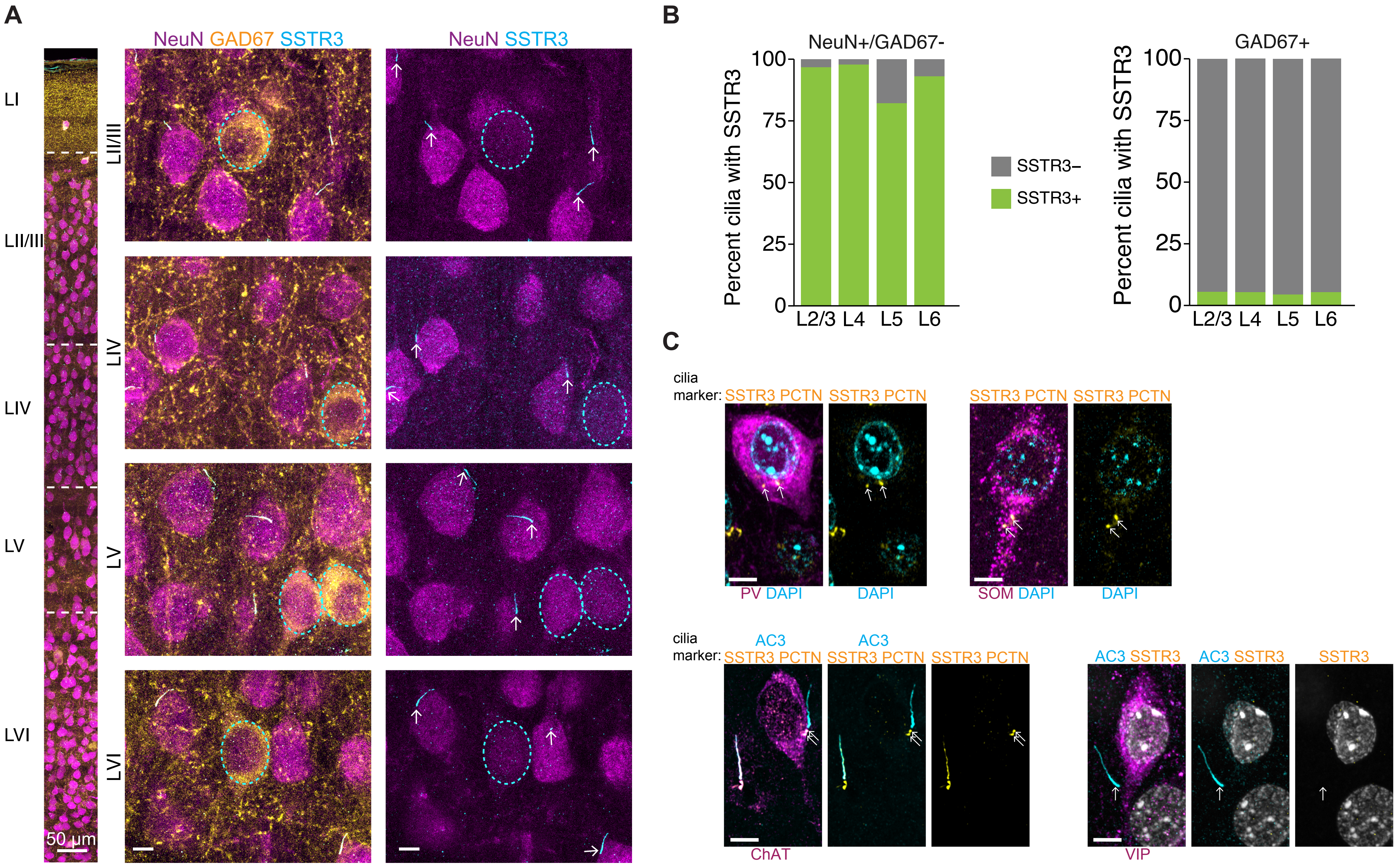


Tereshko et al.,
Figure 5 - figure supplement 1

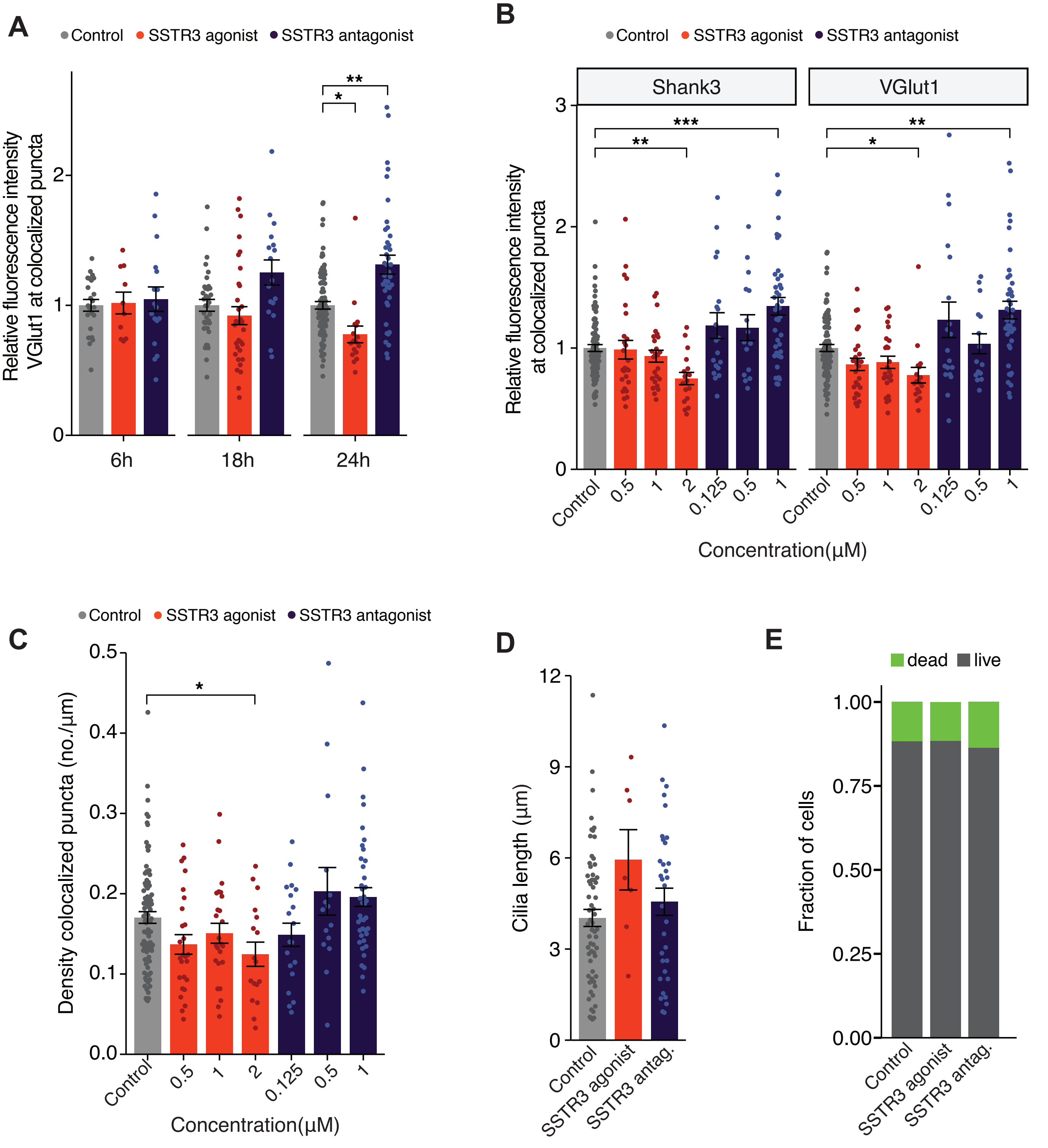


Tereshko et al.,
Figure 6 - figure supplement 1

**Table S1.** Plasmids used in this work.

| Plasmid Name | Description | Source | Catalog or ID # |
|---|---|---|---|
| pLKO.1 | pLKO.1 empty vector | David Root via Addgene | Addgene plasmid #10878 |
| pAAV-hSyn-EGFP | pAAV-hSyn-EGFP | Bryan Roth via Addgene | Addgene plasmid #50465 |
| pSUPER | pSUPER empty vector | OligoEngine | OligoEngine VEC-pBS-0002 |
| pLRT18 | pSUPER-H1-shCep164 | Cloned into pSUPER | OligoEngine VEC-pBS-0002 |
| pLRT19 | pLKO-U6-shArl13b_1 | Cloned into pLKO.1 | Addgene plasmid #10878 |
| pLRT26 | pSUPER-H1-shIft88 | Cloned into pSUPER | OligoEngine VEC-pBS-0002 |
| pLRT67 | pAAV-H1-shArl13b_2-hSyn-EGFP | Modified Addgene plasmid #50465 and pSUPER | Addgene plasmid #50465 |
